# Lark Sparrows (*Chondestes grammacus*) have Ecogeographic Song Variation across North America

**DOI:** 10.64898/2026.08.25.745997

**Authors:** Sofia H Fuertes, Kaiya L Provost

## Abstract

Machine learning models can be used to analyze large bioacoustics datasets and explore variation due to geography or habitat. We find that in the monotypic Lark Sparrow (*Chondestes grammacus*), both environmental variables and geographic distance influence song variation in this species. Bird song is an important method of communication within avian species. The variation in bird song within a species can be due to a variety of factors, including genetic drift and isolation by distance. However, it remains unclear in species with wide ranges how environmental factors in particular can cause changes to the song. In this study, the song *C. grammacus* was analyzed via machine learning to determine if it had significant variation based on multiple geographical metrics. We trained a convolutional neural network to segment individual syllables of 91 *C. grammacus* recordings, then extracted song characteristics. We found that ecoregion and state explain variation in *C. grammacus* songs. Our results demonstrate the efficacy of using machine learning models to analyze large datasets, as well as the impact that ecogeographic variation has on song variance.

**Lay Summary:**

- Birds often have a lot of variation in how they sound depending on where they live, but it is difficult to know what causes this variation without a lot of recordings
- The number of recordings that are needed to understand sound variation at continent-level scales means that analysis by hand is too time consuming
- We used machine learning to process the sound variation which let us understand all Lark Sparrow song in North America
- Lark Sparrows sing differently in different habitat types and they also sing differently when they are far from one another
- This variation is important to understand because it can help these species communicate across wide ranges of environments

## Introduction

Vocalization is important for bird communication (Catchpole and Slater 2008, Brumm and Zollinger 2013), particularly in songbirds, where song is primarily used for reproductive purposes (i.e. to repel intruders and attract mates; Nowicky and Searcy 2006, Kroodsma and Byers 1991). Variations in these vocalizations are often important for recognition between conspecifics (Nelson 1989, Freeberg et al. 1995, Lemon 1966), and even between members of the same population (Dingle et al. 2010), Nowicki and Searcy 2005). As such, geographic variation, often called ‘dialect’ (Baker and Cunningham 1985), can reflect meaningful changes in reproductive behaviors. Geographic variation in bird song occurs in many species and is due to several different factors, both biotic and abiotic. First, the environment can play direct and indirect roles on song (Rodríguez 2002, Derryberry et al. 2009, Brumm and Naguib 2009, Henry and Lucas 2010). Climate, urbanization, humidity, habitat, and tree cover can all affect how song travels, influencing song. Second, like all phenotypes, song is partially genetically determined, meaning that it is subject to isolation-by-distance and genetic drift (Aplin 2019). When a species is separated by large geographic distance, mating cannot occur between separated populations, leading to genetic variation between the two groups. Additionally, random events can cause changes in the genetic composition of a population, changing the predominant traits purely by chance. Finally, unlike other forms of communication in bird species, song is a learned behavior (Slater 1988, Slater 1986, Slater 1989, Slabbekoorn and Smith 2002, Podos and Warren 2007, Derryberry 2009). This means that it is subject to cultural evolution, which is caused by mistakes in learning through time (Aplin 2019). This can lead to song variation across space and time.

To understand geographic variation of song, a thorough quantification of the syllables is needed. A syllable is a single, distinct unit of noise that makes up a bird’s song. Different combinations of syllables create a bird’s unique song, and their layout is known as syntax. In order to analyze a bird’s song, quantification and identification of syllables is needed. However, the quantification of syllables across large geographic regions can be time consuming when done manually, particularly because of the sheer dataset sizes. As such, more automated ways to identify and analyze individual syllables is required. Machine learning is a useful tool that is being applied to biology and bioacoustics, allowing for complex analysis of large datasets (Stowell 2022). This technology can be used to automate syllable detection across many more individuals than was previously feasible by hand. One example of this is TweetyNet, a neural network that is used to automatically annotate the syllables of a birdsong spectrogram (Cohen et al. 2022). TweetyNet has been successfully used by other bioacoustic researchers to accurately segment spectrograms into syllables for a variety of bird species, yielding results with high accuracy (Yang et al. 2023). The variations in volume, frequency, and pattern of bird song allow for quantitative analysis by isolating each individual syllable sung by a bird, and studying their usage in relation to other syllables. The qualities of each bird’s song can then be compared across populations to determine geographical patterns.

In this study we use machine learning to analyse the geographic variation in song of the Lark Sparrow (*Chondestes grammacus*). We chose this species because it is widespread across North America and has songs with strong variation across individuals within the species (e.g., Pandalfino 2019). We hypothesize that the Lark Sparrow’s song is shaped by cultural evolution and environmental context. Thus, we predict to see statistically significant variation in Lark Sparrow song samples due to ecoregion and geography.

## Methods

Generative AI was not used in the production of this manuscript.

### Song Downloads and Annotations

We downloaded 147 recordings of *C. grammacus* from the sound database Xeno-Canto (xeno-canto.org) using the package warbleR version 1.1.30 (Araya-Salas and Smith-Vidaurre, 2017) in R version 4.3.1 (R Core Team, 2023). We only downloaded A, B, and C quality recordings. Of these recordings, 10 were removed from the dataset due to the presence of multiple singing individuals whose songs overlapped, and 46 were removed due to downstream processing issues, leaving 91 (Table 1).

**Table 1:** All Lark Sparrow recordings used for model training and analysis. These recordings were obtained from Xeno-Canto. ID = Xeno-Canto ID number. Lat = latitude. Long = longitude. Q = quality score, with A indicating best quality. Ecoregions are given with the following details: “Mediterranean CA” = Mediterranean California, “E. Temperate Forests” = Eastern Temperate Forests, “S. Semiarid Highlands” = Southern Semiarid Highlands. Recordings with ecoregion listed as “n/a” were excluded from analyses due to data quality issues. Recordings used to train, validate, or test model show the number of seconds used in each data partition. “Train” refers to the seconds of recording used to train the model. “Val” refers to the seconds of recording used to validate the model. “Test” refers to the seconds of recording used to test the model. There was a 10:1:1 split of the data between train:val:test.

| ID | Lat | Long | Q | Ecoregion | State | Train | Val | Test |
| --- | --- | --- | --- | --- | --- | --- | --- | --- |
| XC001257 |  |  | A | n/a | Colorado | 1.95 |  |  |
| XC021386 | 25.97 | -111.92 | A | North American Deserts | Baja California Sur |  |  |  |
| XC031027 | 41.88 | -83.69 | C | E. Temperate Forests | Michigan |  |  |  |
| XC058813 | 32.41 | -94.83 | A | E. Temperate Forests | Texas | 15.92 |  |  |
| XC072198 | 32.04 | -115.91 | B | Mediterranean CA | Baja California Sur |  |  |  |
| XC072199 | 32.04 | -115.91 | A | Mediterranean CA | Baja California Sur | 8.95 |  | 3.43 |
| XC072200 | 32.04 | -115.91 | A | Mediterranean CA | Baja California Sur | 27.16 | 5.88 | 2.35 |
| XC076967 | 39.22 | -108.87 | A | North American Deserts | Colorado | 18.92 | 5.18 |  |
| XC111496 | 31.91 | -109.15 | B | S. Semiarid Highlands | Arizona |  |  |  |
| XC111497 | 31.91 | -109.15 | B | S. Semiarid Highlands | Arizona |  |  |  |
| XC111498 | 31.84 | -109.03 | C | S. Semiarid Highlands | New Mexico |  |  |  |
| XC111499 | 31.84 | -109.03 | C | S. Semiarid Highlands | New Mexico |  |  |  |
| XC111500 | 31.84 | -109.03 | B | S. Semiarid Highlands | New Mexico |  |  |  |
| XC111501 | 31.84 | -109.03 | B | S. Semiarid Highlands | New Mexico |  |  |  |
| XC111502 | 31.84 | -109.03 | B | S. Semiarid Highlands | New Mexico |  |  |  |
| XC111503 | 31.84 | -109.03 | B | S. Semiarid Highlands | New Mexico |  |  |  |
| XC111504 | 31.84 | -109.03 | C | S. Semiarid Highlands | New Mexico |  |  |  |
| XC111505 | 31.84 | -109.03 | B | S. Semiarid Highlands | New Mexico |  |  |  |
| XC111506 | 31.84 | -109.03 | B | S. Semiarid Highlands | New Mexico |  |  |  |
| XC111507 | 31.34 | -109.27 | B | S. Semiarid Highlands | Arizona |  |  |  |
| XC111508 | 31.91 | -109.15 | B | S. Semiarid Highlands | Arizona |  |  |  |
| XC111525 | 31.84 | -109.03 | B | S. Semiarid Highlands | New Mexico |  |  |  |
| XC111527 | 31.84 | -109.03 | B | S. Semiarid Highlands | New Mexico |  |  |  |
| XC111528 | 31.84 | -109.03 | C | S. Semiarid Highlands | New Mexico |  |  |  |
| XC111529 | 31.84 | -109.03 | B | S. Semiarid Highlands | New Mexico |  |  |  |
| XC111530 | 31.84 | -109.03 | B | S. Semiarid Highlands | New Mexico |  |  |  |
| XC111531 | 31.84 | -109.03 | B | S. Semiarid Highlands | New Mexico |  |  |  |
| XC111532 | 31.84 | -109.03 | C | S. Semiarid Highlands | New Mexico |  |  |  |
| XC111533 | 31.34 | -109.27 | C | S. Semiarid Highlands | Arizona |  |  |  |
| XC111534 | 31.34 | -109.27 | C | S. Semiarid Highlands | Arizona |  |  |  |
| XC111535 | 31.34 | -109.27 | C | S. Semiarid Highlands | Arizona |  |  |  |
| XC125423 | 32.98 | -116.60 | A | Mediterranean CA | California | 7.01 |  |  |
| XC125424 | 35.66 | -118.03 | B | North American Deserts | California |  |  |  |
| XC125426 | 32.98 | -116.60 | A | Mediterranean CA | California | 28.98 |  |  |
| XC125427 | 32.98 | -116.60 | A | Mediterranean CA | California | 22.39 |  | 2.21 |
| XC125430 | 34.19 | -116.90 | A | Mediterranean CA | California | 24.57 | 6.52 | 7.79 |
| XC125433 | 34.19 | -116.90 | A | Mediterranean CA | California | 39.46 | 5.45 | 4.61 |
| XC125445 | 34.19 | -116.90 | A | Mediterranean CA | California | 12.87 |  |  |
| XC125447 | 35.66 | -118.03 | A | North American Deserts | California | 13.83 | 1.29 | 12.51 |
| XC131483 | 38.17 | -109.39 | A | North American Deserts | Utah | 18.71 | 2.46 |  |
| XC133060 | 30.08 | -99.50 | A | Great Plains | Texas |  |  |  |
| XC139903 | 41.31 | -105.58 | B | North American Deserts | Wyoming |  |  |  |
| XC141484 | 30.08 | -99.50 | B | Great Plains | Texas |  |  |  |
| XC141485 | 30.08 | -99.50 | B | Great Plains | Texas |  |  |  |
| XC141486 | 30.08 | -99.50 | B | Great Plains | Texas |  |  |  |
| XC141487 | 30.08 | -99.50 | B | Great Plains | Texas |  |  |  |
| XC141488 | 30.08 | -99.50 | B | Great Plains | Texas |  |  |  |
| XC205472 | 38.28 | -108.72 | A | n/a | Colorado | 11.62 | 3.27 | 3.43 |
| XC221053 | 23.42 | -106.25 | A | n/a | Sinaloa | 10.14 | 3.60 |  |
| XC237383 | 39.10 | -93.16 | A | n/a | Missouri | 18.65 | 3.73 |  |
| XC277755 | 38.17 | -98.49 | A | n/a | Kansas | 13.72 |  |  |
| XC277765 | 42.52 | -100.56 | A | n/a | Nebraska | 14.72 |  |  |
| XC323528 | 32.59 | -109.85 | A | n/a | Arizona | 16.13 |  |  |
| XC361883 | 36.68 | -121.21 | A | n/a | California | 6.80 |  |  |
| XC424774 | 37.63 | -104.32 | A | Great Plains | Colorado | 5.33 |  |  |
| XC453098 | 33.03 | -116.95 | B | Mediterranean CA | California |  |  |  |
| XC453920 | 31.99 | -110.96 | C | North American Deserts | Arizona |  |  |  |
| XC461108 | 32.62 | -116.60 | B | Mediterranean CA | California |  |  |  |
| XC462019 | 37.32 | -119.81 | B | Mediterranean CA | California |  |  |  |
| XC465658 | 35.16 | -119.82 | B | Mediterranean CA | California |  |  |  |
| XC483018 | 42.81 | -118.86 | A | North American Deserts | Oregon |  |  |  |
| XC510873 | 41.59 | -118.74 | B | North American Deserts | Nevada |  |  |  |
| XC538538 | 32.67 | -116.93 | A | Mediterranean CA | California |  |  |  |
| XC539282 | 32.67 | -116.93 | A | Mediterranean CA | California |  |  |  |
| XC548571 | 32.67 | -116.93 | A | Mediterranean CA | California |  |  |  |
| XC554528 | 32.67 | -116.93 | B | Mediterranean CA | California |  |  |  |
| XC554578 | 35.13 | -119.84 | C | Mediterranean CA | California |  |  |  |
| XC565246 | 38.76 | -121.68 | A | Mediterranean CA | California |  |  |  |
| XC572845 | 40.58 | -116.56 | B | North American Deserts | Nevada |  |  |  |
| XC596715 | 43.27 | -118.85 | A | North American Deserts | Oregon | 26.05 |  |  |
| XC639148 | 38.53 | -121.09 | B | Mediterranean CA | California |  |  |  |
| XC640973 | 34.54 | -86.80 | B | E. Temperate Forests | Alabama |  |  |  |
| XC660258 | 40.87 | -120.03 | A | North American Deserts | California |  |  |  |
| XC684352 | 32.32 | -100.05 | A | Great Plains | Texas |  |  |  |
| XC695159 | 32.22 | -110.82 | C | North American Deserts | Arizona |  |  |  |
| XC734596 | 41.96 | -83.94 | C | E. Temperate Forests | Michigan |  |  |  |
| XC734597 | 41.96 | -83.94 | C | E. Temperate Forests | Michigan |  |  |  |
| XC741487 | 49.29 | -119.51 | A | North American Deserts | British Columbia |  |  |  |
| XC791866 | 32.67 | -116.93 | A | Mediterranean CA | California |  |  |  |
| XC791979 | 38.52 | -121.10 | B | Mediterranean CA | California |  |  |  |
| XC800855 | 38.41 | -122.08 | A | Mediterranean CA | California |  |  |  |
| XC802849 | 38.44 | -122.12 | A | Mediterranean CA | California | 17.06 | 2.47 |  |
| XC808829 |  |  | A | n/a | California | 12.31 |  | 3.28 |
| XC892364 | 29.24 | -97.29 | A | Great Plains | Texas |  |  |  |
| XC896993 | 32.67 | -116.93 | A | Mediterranean CA | California |  |  |  |
| XC898676 | 39.00 | -122.22 | A | Mediterranean CA | California | 4.23 | 1.03 | 2.27 |
| XC900999 | 25.99 | -97.43 | A | Great Plains | Texas | 2.78 |  |  |
| XC914441 | 24.94 | -99.60 | A | Great Plains | Nuevo Leon |  |  |  |
| XC914442 | 24.94 | -99.60 | A | Great Plains | Nuevo Leon |  |  |  |
| XC914675 | 41.04 | -119.98 | A | North American Deserts | Nevada |  |  |  |

Recordings that were denoted as A quality had each sung syllable annotated by hand using Raven Pro 1.6 (K. Lisa Yang Center for Conservation Bioacoustics 2026). All recordings were analyzed with a spectral window size (i.e. Fast Fourier Transform size) of 256 samples. These annotations separated each individual syllable sung by the Lark Sparrow based on time and frequency.

### TweetyNet

We chose 26 annotated A-quality recordings to train the TweetyNet model (Table 1). The 26 recordings chosen covered the full range of where the Lark Sparrow could be found to make for an unbiased training data set. All syllables were labeled as 1, while noise was labeled as 0. The recordings were converted from mp3 files to wav files, standardized to a 48,000 hz sampling rate because it was the most common among the samples, using the package tuneR version 1.4.7 (Ligges et al. 2023) in R. The annotations were then chopped into smaller files to balance the amount of sound and silence in the recordings. A large expanse of non-annotated silence can cause the model to have an inflated accuracy rate, so those were removed from the recordings (Provost et al. 2022). After chopping, the text files were converted to csv files in simple sequence format using a custom script. The wav and csv files were then used to train TweetyNet version 0.8.0 (Cohen et al. 2022). This was done using the vak library version 1.0.0 (Cohen et al. 2022) in Python version 3.10.14. There was a total of 42 seconds of test data, 400 seconds of training data, and 41 seconds of validation data (Table 1). Once trained, we had the model predict the annotations of data it had never seen before. This model was then used to annotate B and C quality recordings. A total of 65 recordings of B and C quality were annotated this way.

### Song Properties

Summary statistics for all 91 recordings were generated using warbleR on the automatic annotations. Of the 26 recordings that were used to train the TweetyNet model, 10 failed to be processed by the pipeline and were removed from the analysis, leaving 81 recordings to be evaluated. These 81 recordings consisted of 10,805 syllables (Supplemental Table 1).

### Environmental Data

We used multiple metrics to categorize the range across which the Lark Sparrow exists, including latitude/longitude, state, and ecoregion. Ecoregions were used as a proxy for areas with similar environments (Smith et al. 2020) and were sourced from the Commission for Environmental Cooperation (CEC 2021). Recordings were distributed across 14 states/provinces (Canada: British Columbia; USA: California, Nevada, Oregon, Arizona, Utah, New Mexico, Colorado, Wyoming, Texas, Alabama, Michigan; Mexico: Baja California Sur, Nuevo Leon). There were five ecoregions present in our dataset: Mediterranean California, Southern Semiarid Highlands, North American Deserts, Great Plains, and Eastern Temperate forests. Each ecoregion had at least two states associated with it, with North American Deserts having the most states (nine), and five states being associated with multiple ecoregions.

### Analyses

A total of 141 summary statistics were generated. These capture different variations that can be found in bird song, including frequency, bandwidth, duration, amplitude, etc. After removing correlated variables (absolute coefficient correlation >0.75), we kept 19 summary statistics (Table 2; Supplemental Figure 1). We then performed a principal components analysis (PCA) on these 19 variables, which were centered and scaled. We used a broken stick analysis to determine which of our principal components to retain for further analysis, of which we retained five (Peres-Neto et al. 2003; Table 2).

**Table 2:**
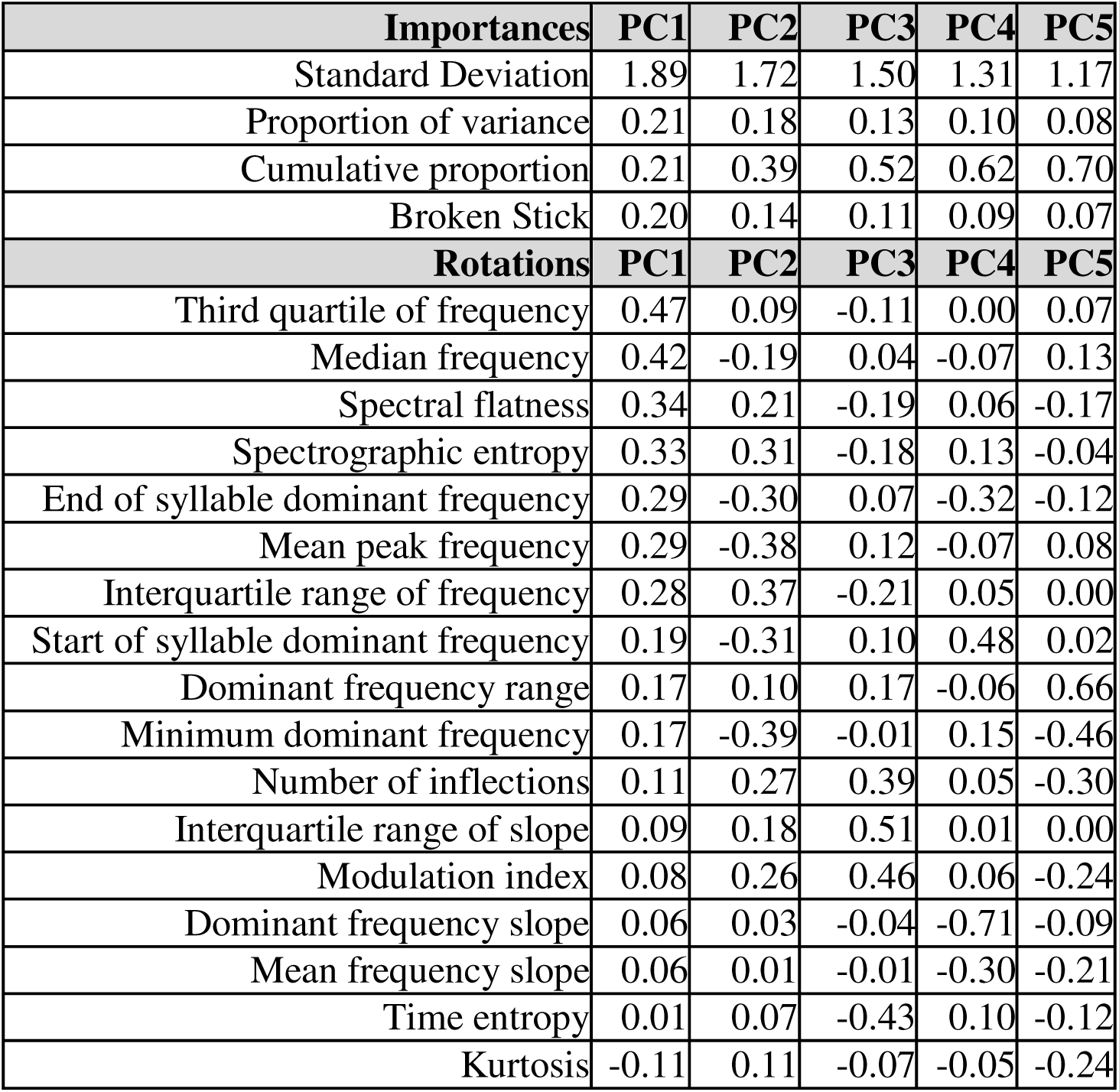
Summary of PCA of Lark Sparrow song. Components 1-5 were retained due to their location below the best fit line of the broken stick model. The proportion of variance indicates the extent to which that PC explains the variance in the data. The cumulative proportion similarly shows the total variance explained by the PC and all previous PCs, showing that the 5 retained principal components explain 70% of all variance.

| **Importances** | **PC1** | **PC2** | **PC3** | **PC4** | **PC5** |
| --- | --- | --- | --- | --- | --- |
| Standard Deviation | 1.89 | 1.72 | 1.50 | 1.31 | 1.17 |
| Proportion of variance | 0.21 | 0.18 | 0.13 | 0.10 | 0.08 |
| Cumulative proportion | 0.21 | 0.39 | 0.52 | 0.62 | 0.70 |
| Broken Stick | 0.20 | 0.14 | 0.11 | 0.09 | 0.07 |
| **Rotations** | **PC1** | **PC2** | **PC3** | **PC4** | **PC5** |
| Third quartile of frequency | 0.47 | 0.09 | -0.11 | 0.00 | 0.07 |
| Median frequency | 0.42 | -0.19 | 0.04 | -0.07 | 0.13 |
| Spectral flatness | 0.34 | 0.21 | -0.19 | 0.06 | -0.17 |
| Spectrographic entropy | 0.33 | 0.31 | -0.18 | 0.13 | -0.04 |
| End of syllable dominant frequency | 0.29 | -0.30 | 0.07 | -0.32 | -0.12 |
| Mean peak frequency | 0.29 | -0.38 | 0.12 | -0.07 | 0.08 |
| Interquartile range of frequency | 0.28 | 0.37 | -0.21 | 0.05 | 0.00 |
| Start of syllable dominant frequency | 0.19 | -0.31 | 0.10 | 0.48 | 0.02 |
| Dominant frequency range | 0.17 | 0.10 | 0.17 | -0.06 | 0.66 |
| Minimum dominant frequency | 0.17 | -0.39 | -0.01 | 0.15 | -0.46 |
| Number of inflections | 0.11 | 0.27 | 0.39 | 0.05 | -0.30 |
| Interquartile range of slope | 0.09 | 0.18 | 0.51 | 0.01 | 0.00 |
| Modulation index | 0.08 | 0.26 | 0.46 | 0.06 | -0.24 |
| Dominant frequency slope | 0.06 | 0.03 | -0.04 | -0.71 | -0.09 |
| Mean frequency slope | 0.06 | 0.01 | -0.01 | -0.30 | -0.21 |
| Time entropy | 0.01 | 0.07 | -0.43 | 0.10 | -0.12 |
| Kurtosis | -0.11 | 0.11 | -0.07 | -0.05 | -0.24 |

To investigate the impact of environment and geography on song, 16 linear mixed effects models were made, for each PC, to analyze our data, using the lme4 version 2.0-1 package in R (Bates et al. 2015). A mixed effects model was used to account for the variation due to some syllables being from the same recording, which would lead to similarities in background noise, microphone quality, etc. We treated Recording ID as a random effect. Ecoregion, State, Latitude, and Longitude were treated as fixed effects. All sixteen combinations of these four fixed effects were modelled. The corrected Akaike Information Criterion (AICc) was used to choose the best linear mixed models to describe our data (Burnham and Anderson 2002) using the AICcmodavg version 2.3-4 package in R (Mazerolle 2023). Models with a ΔAICc less than 2.0 were treated as equivalently good.

After identifying the predictors present in the best models (see Results), we then performed post-hoc ANOVAs on categorical predictor variables (Ecoregion, State), including interaction terms if applicable. To evaluate significance, we also performed a Tukey’s Honest Significant Differences test (Yandell 1997). Because we performed four tests, a Bonferroni correction was performed to set the p-value to 0.0125 for significance (0.05/4) (Bonferroni 1936).

## Results

### Machine Learning Model Results

We used TweetyNet to train a model to auto-annotate syllables of the Lark Sparrow. The final model had an accuracy rate of 85.1% on the held out test data.

### Principal Components Analysis

For our principal components analysis, we retained PC1, PC2, PC3, PC4, and PC5 following a broken stick analysis, where these PCs explained more variation than expected. These first five PCs account for 21%, 18%, 13%, 10%, and 8% of the data, respectively (70% total variance).

The PCA rotations showed that PC1 was positively correlated with third quartile frequency, the median frequency, and spectral flatness, and entropy, such that syllables with higher PC1 were overall higher pitched and noisier. PC2 was negatively correlated with mean peak frequency, minimum dominant frequency, and positively correlated with frequency IQR, such that syllables with higher PC2 had lower frequencies and higher bandwidths. PC3 was positively correlated with time entropy and negatively correlated with the number of inflections, slope IQR, and modulation index, such that syllables with higher PC3 were less variable. PC4 was highly negatively correlated with the dominant frequency slope, such that syllables with high PC4 had negative slopes. Lastly, PC5 was positively correlated with dominant frequency range and negatively correlated with minimum dominant frequency, such that syllables with high values of PC5 had low frequencies and high bandwidths. In sum, PC1 reflects noisiness, PC2 and PC5 reflect frequency and bandwidth, PC3 reflects modulation, and PC4 reflects slope.

### Statistical Models

With our sixteen linear mixed models for each PC, we found that overwhelmingly Ecoregion and/or State were the variables present in the best models (Table 3). For PC1 and PC2, the best model contained both Ecoregion and State. PC2 also had an equivalent model (ΔAICc = 1.63) that contained only State. For PC3 and PC4, the best model contained only the random effect of Recording ID, though for PC3, an equivalent model (ΔAICc = 1.33) contained only State. Lastly, for PC5, the best model contained only Ecoregion. Latitude and Longitude were never among the best models (but see Supplemental Table 1).

**Table 3:** Table for mixed effects models for all of them. Bold means equivalently explanatory. All models dAICc less than 2, plus best models with dAICc over 2, shown

| Response | Random Effects | Fixed Effects | AICc | dAICc |
| --- | --- | --- | --- | --- |
| PC1 | Recording ID | Ecoregion, State | 38919.16 | **0.00** |
|  | Recording ID | State | 38921.40 | 2.24 |
| PC2 | Recording ID | Ecoregion, State | 39150.24 | **0.00** |
|  | Recording ID | State | 39151.87 | **1.63** |
|  | Recording ID | Ecoregion, State, Longitude | 39155.56 | 5.32 |
| PC3 | Recording ID | n/a | 38901.04 | **0.00** |
|  | Recording ID | State | 38902.37 | **1.33** |
|  | Recording ID | Ecoregion | 38903.29 | 2.25 |
| PC4 | Recording ID | n/a | 36216.10 | **0.00** |
|  | Recording ID | Longitude | 36225.37 | 9.27 |
| PC5 | Recording ID | Ecoregion | 33025.09 | **0.00** |
|  | Recording ID | Ecoregion, State | 33029.19 | 4.10 |

Because only categorical variables were among the best predictors, we only performed post-hoc ANOVAs on Ecoregion and State. PC4 was excluded as the model with the random effect performed best. For PC1 and PC2 our statistical tests included Ecoregion, State, and their interaction. For PC3 the test only included State, and for PC5 the test only included Ecoregion. When examining the interaction term, we focused specifically on the recordings that were found in the same state but different ecoregion (California, Baja California Sur, Arizona, Colorado, Texas).

For every PC, all terms in the ANOVA were significant (Table 4). Tukey’s HSD tests showed that all ecoregions were significantly in PC1 (*p*<0.003), PC2 (*p*<0.0003), and PC5 (*p*<0.003) with a few exceptions. For PC1, North American Deserts-Mediterranean California and Great Plains-Southern Semiarid Highlands were not significantly different. (*p*>0.9857). For PC2, Eastern Temperate Forests were not significantly different from North American Deserts or Southern Semiarid Highlands (*p*>0.016). For PC5, Southern Semiarid Highlands were not significantly different from North American Deserts or Eastern Temperate Forests (*p*>0.016).

**Table 4:** ANOVA relationship between PC1-PC5 and State or Ecoregion. The predictor “State:Ecoregion” indicates the interaction term between State and Ecoregion. Only recordings that were found in the same state but different ecoregion were considered for the interaction term. All *p* values are significant.

| Response | Predictor | df | Sum Sq | Mean Sq | F | *p* |
| --- | --- | --- | --- | --- | --- | --- |
| PC1 | Ecoregion | 4 | 1457 | 364.2 | 131.6 | 2.0x10<sup>-16</sup> |
|  | State | 13 | 4668 | 359.1 | 129.8 | 2.0x10<sup>-16</sup> |
|  | Ecoregion:State | 1 | 1116 | 1115.9 | 403.3 | 2.0x10<sup>-16</sup> |
| PC2 | Ecoregion | 4 | 1066 | 266.5 | 100.21 | 2.0x10<sup>-16</sup> |
|  | State | 13 | 1911 | 147 | 55.27 | 2.0x10<sup>-16</sup> |
|  | Ecoregion:State | 1 | 579 | 579.4 | 217.84 | 2.0x10<sup>-16</sup> |
| PC3 | State | 13 | 975 | 75.02 | 31.14 | 2.0x10<sup>-16</sup> |
| PC5 | Ecoregion | 4 | 200 | 49.95 | 29.19 | 2.0x10<sup>-16</sup> |

For the states comparisons, all states were significantly different from at least one other state in PC1 (*p*<0.0125), PC2 (*p*<0.011), and PC3 (*p*<0.008) from at least one other state, with the exception of Nuevo Leon, which did not significantly differ from any other state for PC3 (*p*>0.26). Notably, Wyoming was significantly different from every other state in PC1 (*p*<0.0001).

Baja California Sur recordings were significantly different between MC and NAD for both PC1 and PC2 (*p*<0.0001). Arizona recordings were significantly different between NAD and SSH for both PC1 and PC2 (*p*<0.015). Texas recordings were significantly different between GP and ETF for both PC1 and PC2 (*p*<0.0002). Finally, Colorado recordings were significantly different between NAD and GP for only PC2 (*p*<0.0001). California did not show significant differences between NAD and MC (*p*>0.19).

## Discussion

Our retained principal components described 70% of all variance in our data, with PC 1 alone describing 21%. Our principal components analysis shows that this difference is most explained by noisiness, specifically frequency. This shows that the pitch of the Lark Sparrow’s song is the aspect most varied. Variations in frequency have been shown to affect mating success in bird communication (Weisman 2004). It is also shown that pitch can be impacted by habitat noise (Hao et al. 2024).

Latitude and longitude were not significantly different indicators of variation. This indicates that song was not varied across a continuous range, but rather categorically, indicating that those regions have a greater correlation with song variation than simply distance.

Ecoregion was a significant indicator of song variability, supporting our hypothesis that environment would be correlated with variation in birdsong. However, state was the strongest explanatory variable across our five principal components. When comparing across different states within the same ecoregion, variation could still be seen, such as that between Colorado and Baja Sur in North American Deserts (Figure 4). This demonstrates that state lines explain variation even when environments are similar.

**Figure 1:**
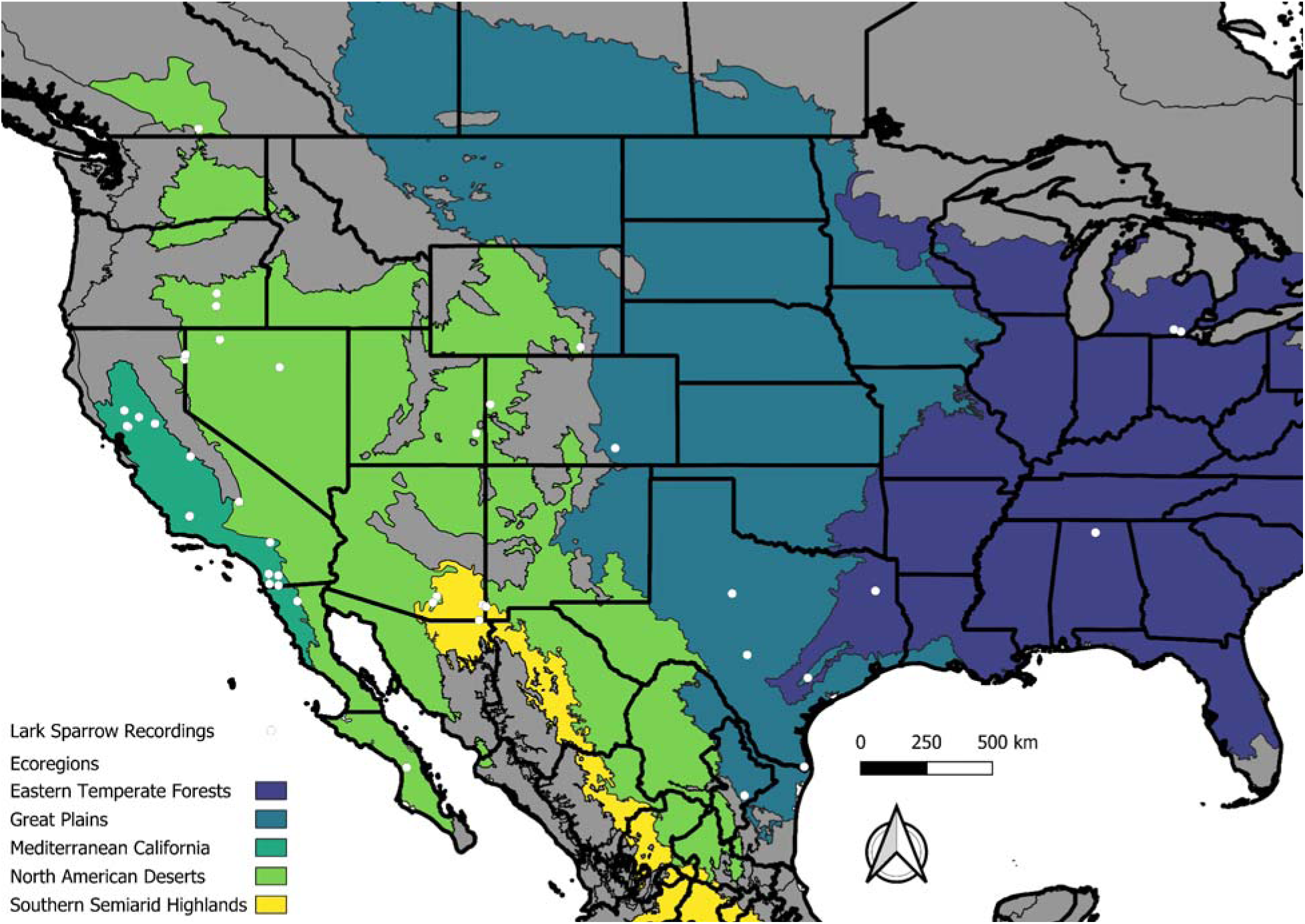
Map showing geographical range of Lark Sparrow recordings across North America. Thick black lines indicate states and provinces. White dots show locations of each recording. Note that multiple recordings may be represented by the same point. Thin black lines and colorful backgrounds indicate ecoregions (navy = Eastern Temperate Forests, teal = Great Plains, dark green = Mediterranean California, light green = North American Deserts, yellow = Southern Semiarid Highlands). Arrow points to true north. Scale bar shows distance in kilometers. These maps were created with QGIS version 3.36.3 (QGIS Development Team, 2024).

**Figure 2:**
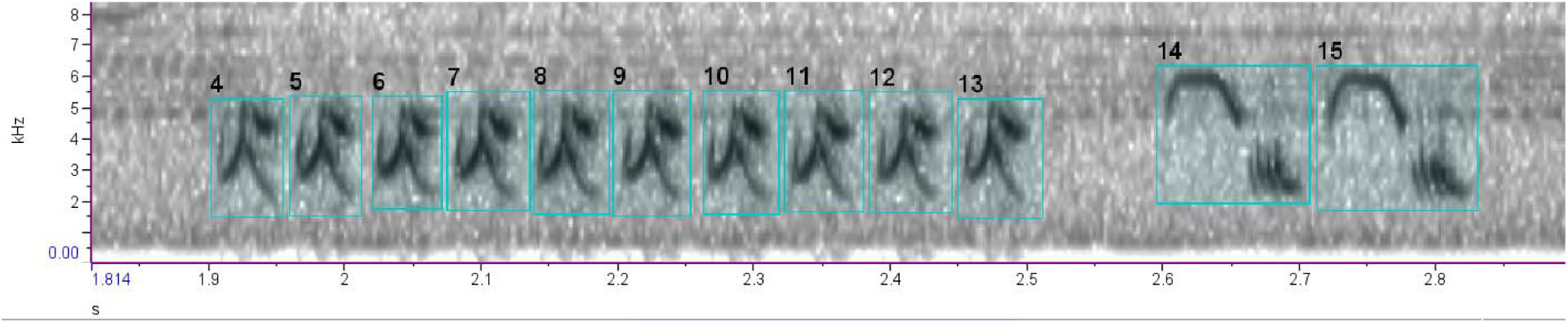
Annotated spectrogram of Lark Sparrow recording XC1257 in Raven Pro 1.6. The x axis is the time in seconds and the y axis is the frequency in kHz. Color indicates amplitude, with higher amplitude frequencies in darker colors. Syllables are numbered from left to right with no overlap. Syllables 4 through 13 are one syllable type, while syllables 14 and 15 are a distinct separate syllable type.

**Figure 3:**
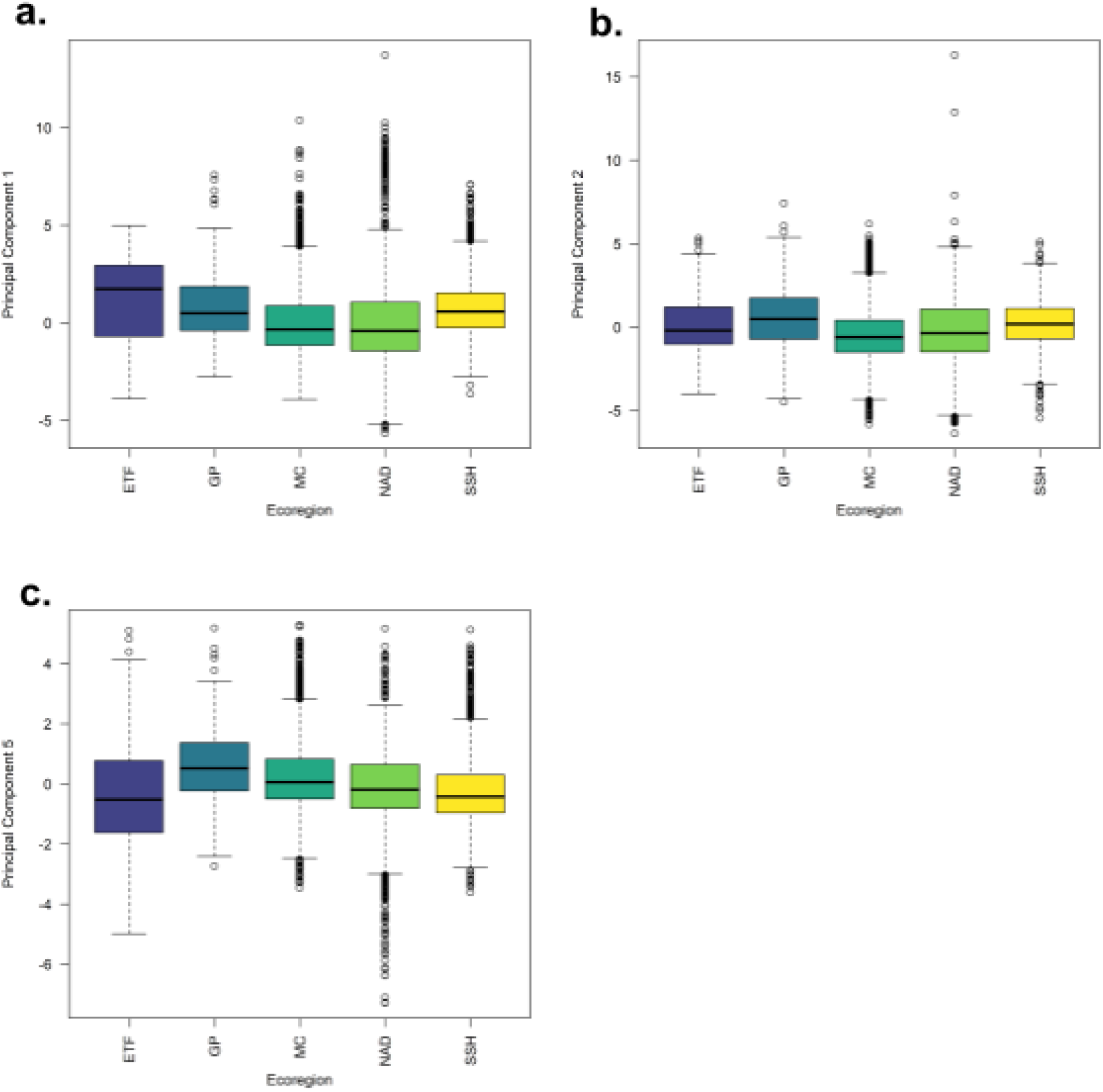
Box plots **a** through **c** show the distribution of principal component scores across the 5 ecoregions represented in our samples. (a) shows PC1. (b) shows PC2. (c) shows PC5. The x axis represents ecoregions, where ETF = Eastern Temperate Forests, GP = great plains, MC = Mediterranean California, NAD = North American Deserts, SSH = Southern Semi-Arid Highlands. Colors of ecoregions correspond to colors in Figure 1.

**Figure 4:** Box plots a and b show the distribution of principal component scores across states that contain multiple different ecoregions (California, Baja California Sur, Arizona, Colorado, Texas). (a) shows PC1. (b) shows PC2. The x axis shows state and ecoregion, where ETF = Eastern Temperate Forests, GP = great plains, MC = Mediterranean California, NAD = North American Deserts, SSH = Southern Semi-Arid Highlands. Colors of ecoregions correspond to colors in Figure 1.

These results support the claim that there is a statistically significant variation in *C. grammacus* song across state and ecoregion, which shows that geography is correlated with vocal variation. The exceptions in statistical significance for Tukey’s HSD test were not similar across PCs, showing that although some regions may share similarities in bird song in one aspect, they may differ in another.

These patterns could be explained by a number of reasons, including cultural evolution, isolation by distance, and genetic drift. These are likely in *C. grammacus* because song is a learned behavior, so small changes can eventually develop into regional dialects over a wide enough range. Analysis of syllabic structure and syntax could further elucidate the specific differences in song between individuals.

Natural selection can also play a major role in this, as different environments can select for different song traits, thus influencing reproductive success (Podos 2022). This matters because of the significant differences in habitat across North America. Ecoregions are defined by their similar environments, meaning that each have distinct climates and ecologies. Differences in humidity, tree cover, weather, and presence of other species can all influence how sound travels and thus influence birdsong (Luther 2009). Additionally, changes in these environments such as global warming, deforestation, or urbanization could have potential effects on bird song, as has been demonstrated in other species (Nemeth 2009). Song plays a significant role in reproductive success, so these changes could have further effects on bird populations and their communities (Kroodsma 1991).

Our machine learning model served as an effective way to analyze a large dataset of bird song recordings across a very wide scope. The model can potentially also be used for birdsong similar to that of the Lark Sparrow. The *Chondestes* genus is monotypic, but it is closely related to the lark bunting (*Calamospiza melanocorys*). Using the same system to annotate the song of *C. melanocorys* could prove useful in determining the model’s versatility.

The recordings range in date, from 1995 to 2024, and the recordings are not evenly distributed throughout that time frame. The distribution of locations over time was not controlled for either. It is possible that there have been changes over this period of time which were not accounted for. Future research can be done on more specific ecoregional variations, such as syllable patterns. *C. grammacus* has a very complex song so it would be worthwhile to see how syntax changes. One could also analyze across multiple species of birds that exist in many different ecoregions and compare song similarity within the ecoregion.

## Acknowledgements

We would like to thank Provost lab members, past and present (Shannon Hong, Ellie Wang, Julissa Flores, Megan Tan, Valeria Lopez, Joseph O’Brien, Annmarie Schurr, Karishma Suchit, Mariah Gordon, Nadiya Boodoo, Tamana Bashardost, June Cumento, Vivian Chow, Skylar Kiang), for their continued support. We thank all reviewers anonymous or otherwise for their feedback. We thank the individuals who have contributed recordings to the Xeno-Canto repository that were used in this study.

## Funding Statement

No additional funding was used to perform this study.

## Ethics Statement

All work was performed following the Adelphi University Code of Ethics.

## Author Contributions

S.H.F. and K.L.P. conceived the idea, design, and experiment. S.H.F. collected data, conducted the research, and analyzed the data. S.H.F. and K.L.P. wrote and edited the paper.

## Conflicts of Interest

The authors of this paper have no conflicts of interest to declare.

## Data Availability

The analyses performed in this article can be reproduced with supplemental data.

## Supplemental Material

**Supplemental Figure 1:**
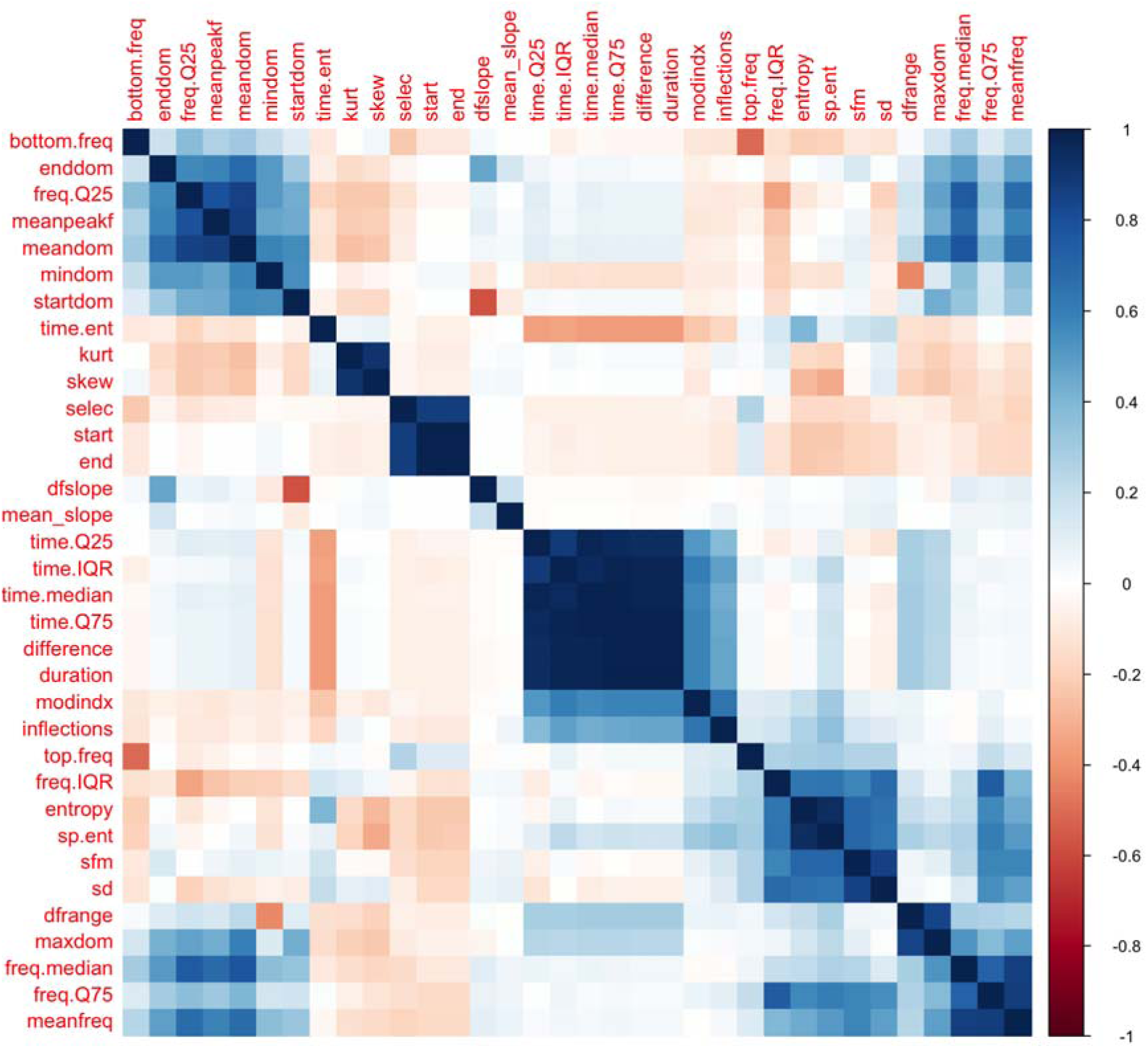
Correlations between song variables. Variables were thinned so that all remaining variables had correlation coefficients -0.75 < r < 0.75. Darker colors indicate stronger correlations. More blue colors indicate positive correlation, more red colors indicate negative correlation, and white indicates no correlation.

**Supplemental Table 1:** Summary of ∼10,000 individual syllables. Attached. Frequency values are in kHz and time values are in seconds unless otherwise specified. Columns designate the following: sound.files: file name of recording; selec: selection in the file; View: whether Spectrogram or Waveforms were used; Channel: Which channel of the recording was used; start, end: Start and ending time in of bounding box; bottom.freq, top.freq: Bottom and top frequency of bounding box; dfrange, dfslope, enddom, startdom, mindom: Dominant frequency range, slope, ending value, starting value, and minimum; entropy: spectrographic entropy; freq.IQR, freq.median, freq.Q75: frequency interquartile range, median, and third quartile value; inflections: number of frequency inflection points; kurt: kurtosis of frequency spectrum ; mean_slope, meanpeakf: mean frequency slope, peak ; modindx: frequency modulation index; sfm: spectral flatness ; time.ent, time.IQR: time entropy and interquartile range; PC1, PC2, PC3, PC4, PC5: principal components 1-5 for sound properties ; Recording_ID: Xeno-Canto recording identification number; Genus, Specific_epithet, Subspecies, English_name: scientific name and English common name information; Recordist: name of individual who recorded Xeno-Canto recording; Country, Locality, Latitude, Longitude: geographic locality information ; Vocalization_type: whether vocalization was a song, call, etc; Quality: ranking of recording quality from A (best) to E (worst); Time, Date: when recording was made ; group: designates recording as part of “birds” collection on Xeno-Canto; Altitude: the altitude of the recording, in meters; sex: sex of the individual recorded, if known; stage: life stage of the individual recorded, if known; ecoregion: ecoregion that recording was made in; State: state or province in which recording was made.

**Supplemental Table 2:**
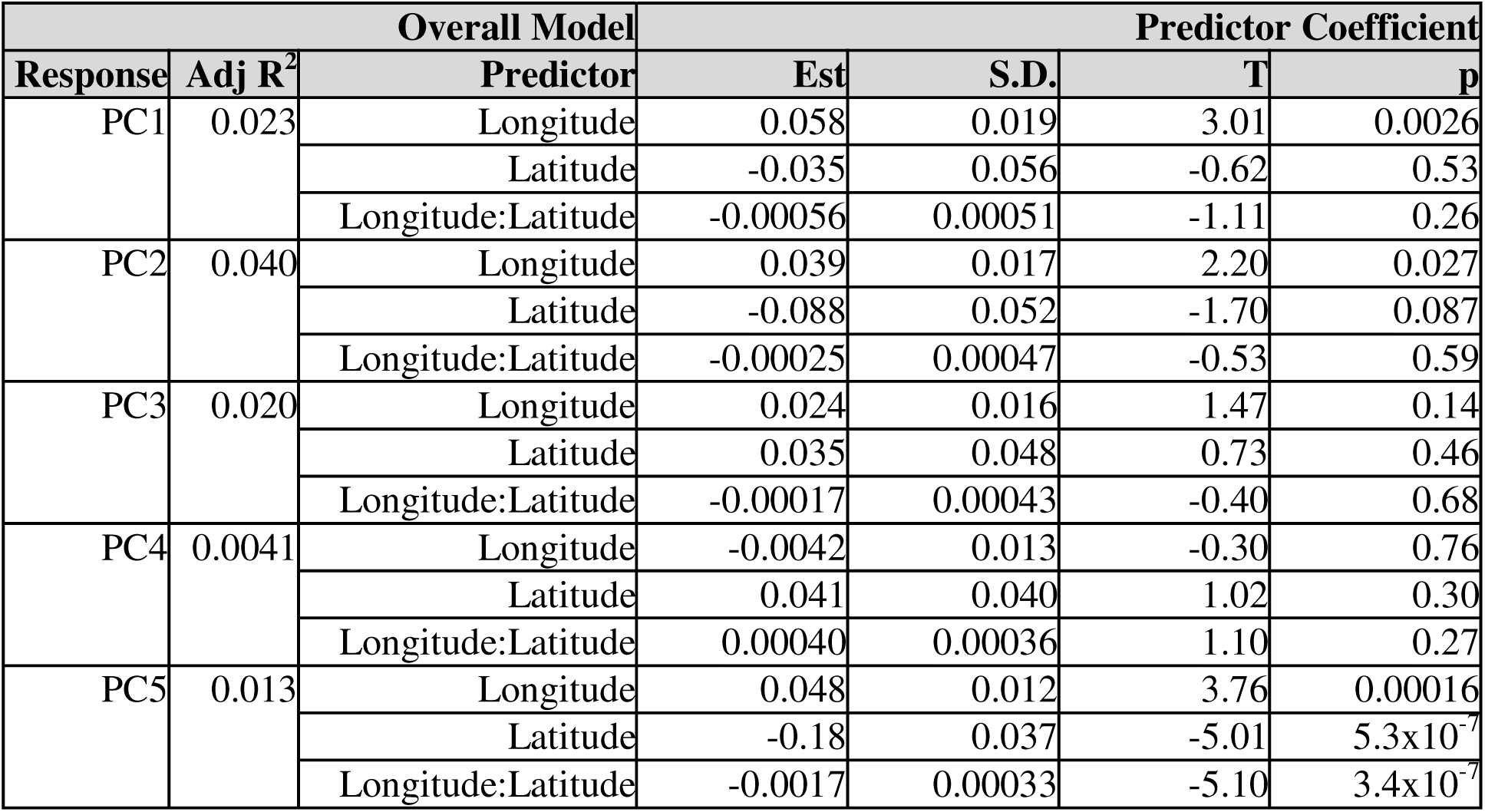
Mixed effect linear model results comparing Principal Components 1-5 with latitude, longitude, and their interaction longitude:latitude. Significant p-values are indicated with bold.

## Notes

### Competing Interest Statement

The authors have declared no competing interest.

